# Biophysically Realistic Neuron Models for Simulation of Cortical Stimulation

**DOI:** 10.1101/328534

**Authors:** Aman S. Aberra, Angel V. Peterchev, Warren M. Grill

## Abstract

1.

**Objective:** We implemented computational models of human and rat cortical neurons for simulating the neural response to cortical stimulation with electromagnetic fields.

**Approach:** We adapted model neurons from the library of Blue Brain models to reflect biophysical and geometric properties of both adult rat and human cortical neurons and coupled the model neurons to exogenous electric fields (E-fields). The models included 3D reconstructed axonal and dendritic arbors, experimentally-validated electrophysiological behaviors, and multiple, morphological variants within cell types. Using these models, we characterized the single-cell responses to intracortical microstimulation (ICMS) and uniform E-field with dc as well as pulsed currents.

**Main results:** The strength-duration and current-distance characteristics of the model neurons to ICMS agreed with published experimental results, as did the subthreshold polarization of cell bodies and axon terminals by uniform dc E-fields. For all forms of stimulation, the lowest threshold elements were terminals of the axon collaterals, and the dependence of threshold and polarization on spatial and temporal stimulation parameters was strongly affected by morphological features of the axonal arbor, including myelination, diameter, and branching.

**Significance:** These results provide key insights into the mechanisms of cortical stimulation. The presented models can be used to study various cortical stimulation modalities while incorporating detailed spatial and temporal features of the applied E-field.

## 2. Introduction

An array of powerful techniques is available for modulating neural activity in the cortex with electromagnetic fields both to understand brain function and to treat neurological as well as psychiatric disorders. Electrical current can be delivered to the brain using intracortical microstimulation (ICMS), epi- and sub-dural stimulation, transcranial electrical stimulation (tES) with either direct (tDCS) or alternating current (tACS), or transcranial magnetic stimulation (TMS), which uses magnetic induction to generate current non-invasively in the brain. However, the effectiveness of these techniques to treat disorders or to probe brain-behavior relationships is hampered by ambiguity and debate about their mechanisms at the cellular level. The cerebral cortex contains densely interwoven cell bodies, axons, and dendrites with incredible diversity of morphological, electrophysiological, and functional characteristics. Using experimental approaches alone, it is challenging to determine which neural elements and populations are modulated by stimulation with specific spatial and temporal parameters. For example, calcium imaging during ICMS with low-amplitude pulse trains suggested that sparse, distributed clusters of neurons are activated [1], while others argue that ICMS can precisely activate local neurons [2]. Epi- and sub-dural stimulation and especially TMS and tES produce much broader electric field (E-field) distributions, and are typically less amenable to simultaneous direct neural recordings, leading to even greater uncertainty about the affected neural populations and their relative thresholds for activation [3].

Computational modeling offers a complementary approach for studying the neural mechanisms of electrical stimulation; models can be used to explore efficiently the very large parameter space, to generate experimentally testable predictions, and to optimize parameters and potentially improve clinical outcomes. Experiments and models show that neural polarization is determined not only by the E-field distribution, but is also highly dependent on neuronal morphology, location, and orientation, particularly of the axons and their collaterals [4–10]. Additionally, generation of APs depends on both passive membrane properties, e.g. capacitance and leak conductance, and active membrane properties, e.g. ion channel densities and kinetics [11,12]. Combining these complex factors in biophysical models that also account for the coupling of the exogenous E-field to the neuronal membranes is critical for accurate predictions of the neural response.

Recent computational studies of electrical stimulation of cortical neurons simulated multi-compartmental neuron models with increasingly realistic morphologies. High-quality, 3D reconstructions of neuronal morphologies can be obtained using software-based tracing of neurons stained with intracellular dyes [13]. However, because of methodological difficulties in obtaining reconstructions of axons [14], most modeling studies use simplified, straight axons [5,15–19]. 3D geometries of intracortical axonal arbors are particularly important for accurate coupling to E-fields. Several studies modeled artificially-defined, branched axons [7,20,21] or white-matter axons defined using tractography based on diffusion tensor imaging [22]. Two studies modeled uniform E-field stimulation of pyramidal cells with reconstructed axon arbors, but used either passive membrane properties [9] or active membrane properties drawn from another model and flattened, 2D representation of the axons [8]. Additionally, the morphological reconstructions were drawn from juvenile rats, and no adjustments were made to account for geometric differences across species or age. Neuronal morphology and physiology differ between species [23,24] and stages of development [25–27]; these factors must be considered in biophysical models of neural stimulation to obtain accurate estimates of the neural response in humans. Eyal et al. used human biopsy samples from temporal cortex to build morphologically-realistic neuron models with membrane and synaptic properties constrained by *in vitro* slice recordings; however, these models lacked axonal arbors beyond the initial segment, making them unsuitable for modeling electrical stimulation [28]. No study has simulated electrical stimulation of models capturing the morphological and functional variations within and between cell types, including validated electrophysiological properties and 3D axonal reconstructions. Therefore, the goal of this study was to implement a library of cortical neuron models to simulate more accurately the direct neural response to electromagnetic stimulation of cortical tissue.

We modified recently published cortical neuron models from the Blue Brain Project [29,30] to reflect the geometrical and biophysical properties of adult rat and human cortical neurons and characterized their response to various forms of cortical stimulation. The cortical neuron models included realistic 3D axon morphologies, experimentally validated electrophysiological responses, and morphological diversity both within and across excitatory as well as inhibitory cell types. The models reproduced experimental data from ICMS as well as uniform E-field stimulation, and confirmed that the direct response to stimulation is highly dependent on axonal morphology, including diameter, orientation, and distribution of terminal branches, as well as the presence of myelination. This work provides biophysically-realistic compartmental neuron models for studying the mechanisms of different brain stimulation modalities.

## 3. Methods

We adapted models of cortical neurons with realistic axonal and dendritic morphologies for simulations of the direct, single-cell response to cortical stimulation using electric fields generated by either current injection or magnetic induction. We studied both suprathreshold stimulation with ICMS using non-uniform E-fields generated by a microelectrode, as well as suprathreshold stimulation and subthreshold polarization using a uniform E-field, similar to the local field generated by transcranial electric and magnetic stimulation.

### 3.1 Blue Brain Model Neurons

The neural models were modified versions of the multi-compartmental, conductance-based models implemented by the Blue Brain Project [29,30] in the NEURON v7.4 simulation software [31]. The original model morphologies were obtained from 3D digital reconstructions of biocytin-filled neurons in all 6 cortical layers of slices of somatosensory cortex from P14 male Wistar rats. To increase morphological diversity amongst cells of the same type, Markram et al. generated virtual “clones” by adding random variation to the branch lengths and rotations of the exemplar models [29]. A feature-based multi-objective optimization method [32] was used to fit to electrophysiological data the conductances of up to 13 different published Hodgkin-Huxley-like ion channel models in soma, basal dendrites, apical dendrites, and axon initial segment. The ion channels included transient sodium, persistent sodium, transient potassium, persistent potassium, M-current (Kv7), H-current, high-voltage-activated calcium, low-voltage-activated calcium, A-type potassium (Kv3.1), D-type potassium (Kv1), stochastic potassium, and SK calcium-activated potassium [29]. From this library of 207 cell types, we selected cells based on their relative abundance and refined our selection within each layer based on cell types exhibiting the lowest thresholds for stimulation with extracellular electric fields. The final set of cell types included layer 1 (L1) neurogliaform cell with a dense axonal arbor (NGC-DA), L2/3 pyramidal cell (L2/3 PC), L4 large basket cell (L4 LBC), L5 thick-tufted pyramidal cell with an early bifurcating apical tuft (L5 TTPC), and L6 tufted pyramidal cell with its dendritic tuft terminating in L4 (L6 TPC-L4). Figure 1 depicts the morphologies of the adult rat model neurons, and the adult human versions are included in the supplemental materials (Figure S1).

**Figure 1.**
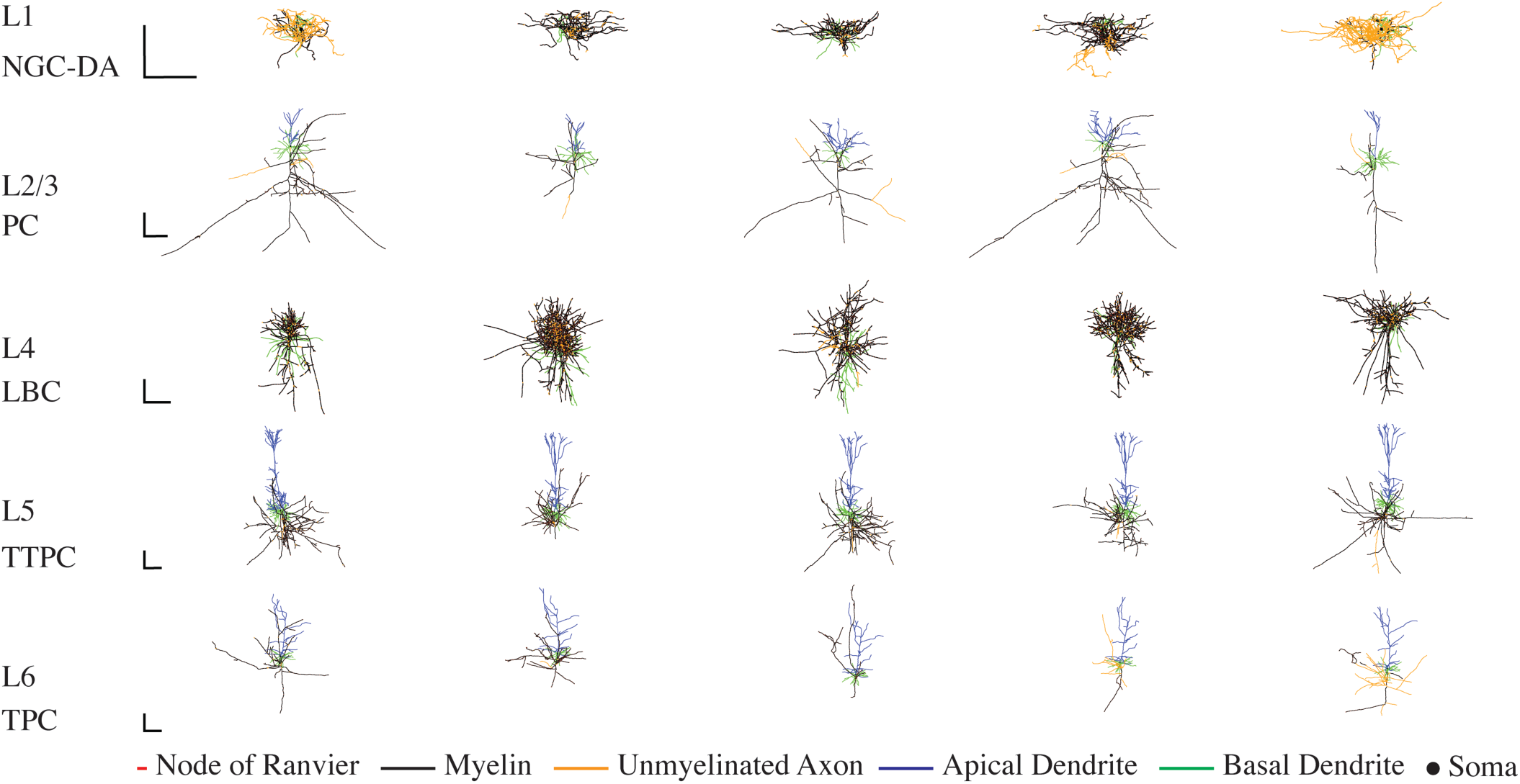
Morphology of model cortical neurons. 3D reconstructions of morphological cell types used in this study. Each row contains the 5 clones of each cell type drawn from the layer indicated on the left. Cell type abbreviations defined in Section 3.1. Adult rat versions shown here, with morphologies colored to indicate nodes of Ranvier (red), myelin (black), and unmyelinated axonal sections (yellow), as well as apical dendrites (blue) and basal dendrites (green). Scale bars = 250 *μ*m.

Markram et al. used automated “repair” algorithms to regrow computationally axons that were cut by the slice sectioning, thereby generating axonal morphologies that matched the statistics of intact axons for each neuron model [14]. The full axonal arbors were used to estimate synaptic connectivity statistics but were not included in the parameter optimizations or network simulations in the original study [29]. Since axons are likely the lowest threshold elements for most forms of electrical stimulation [12,33,34], these axonal arbors were included in our models after the modifications described below. The NEURON code to generate these modified neuron models is available on ModelDB (https://senselab.med.yale.edu/modeldb/ShowModel.cshtml?model=241165).

### 3.2 Modifications to Blue Brain Model Neurons

#### 3.2.1 Myelination

The vast majority of intracortical axons have some myelination; the extent and pattern of myelin coverage varies based on a number of factors, including cell type, layer, brain-region, and species [35–39]. The axon collaterals of neurons in motor and somatosensory cortices are also myelinated [37,40], particularly their horizontal branches, before dividing into finer, unmyelinated processes [41,42]. Myelinated axons have shorter chronaxies and lower thresholds for activation by electrical stimulation [12], making them likely targets for activation with short pulse durations, e.g., ICMS and TMS.

The axons in the Blue Brain models were unmyelinated; therefore, an automatic myelination algorithm was implemented to approximate myelination of adult neocortical axonal arbors while preserving the original reconstructed morphologies (Figure 2a). This algorithm registered nodes of Ranvier to bifurcations [43–45], with nodal lengths of 1 *μ*m [46], and replaced the intermediate branches with internodal myelinated sections and nodes of Ranvier. The internodal lengths (*L*) were calculated from the diameter of the original compartments based on the ratio of internodal lengths to fiber diameter (*D*) with *L/D*=100 [47–49]. The ratio of the outer myelin diameter to the inner axon diameter was varied as a function of inner diameter by fitting third order polynomials to measurements from [35]. For pre-terminal branches, the ratio of internodal length to fiber diameter was changed to 70, to approximate the shorter myelinated segments observed near terminals [44]. Terminals were modeled with the same geometric properties as nodes of Ranvier. Axonal branches shorter than 20 *μ*m or thinner than 0.2 *μ*m were left unmyelinated [46,50].

**Figure 2.**
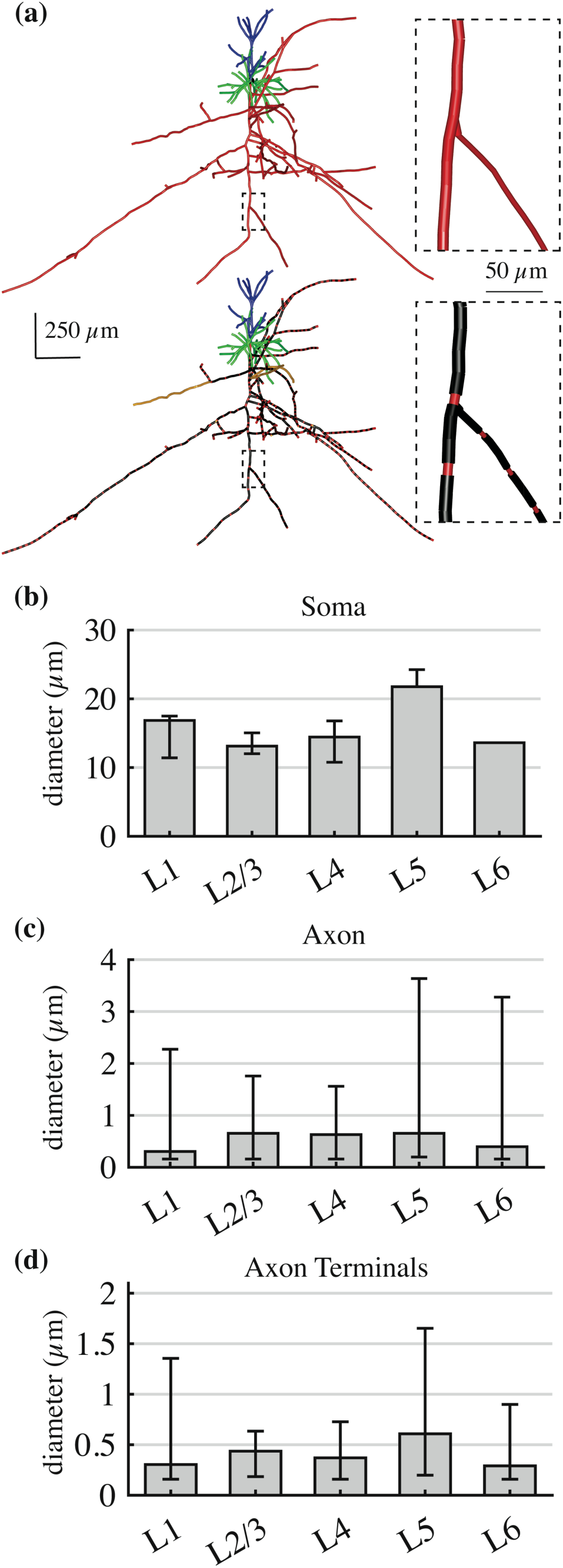
Modifications to approximate adult rat cortical neurons. (**a)** Myelinated axonal branches in example L2/3 PC. The myelination algorithm preserved the original geometry, but replaced existing sections with myelinated internodal sections and nodal sections according to the axon diameter. (**b)** – (**d**) Median diameter (± minimum/maximum across 5 clones within cell type) of **(b)** soma major axis, (**c)** all axonal compartments, and (**d)** axon terminal compartments.

#### 3.2.2 Myelinated Axon Biophysics

To approximate axonal biophysics in each cell type with minimal assumptions or deviations from the originally optimized models, nodal sections were modeled with the same ion channels present in the axon initial segment, except the calcium channels were removed [51] and the transient sodium channel conductance, specific to each cell model’s initial segment, was doubled, based on the density of Nav1.6 channels [52] (Table 1). Unmyelinated branches had the same sodium channel density as the initial segment.

**Table 1.**
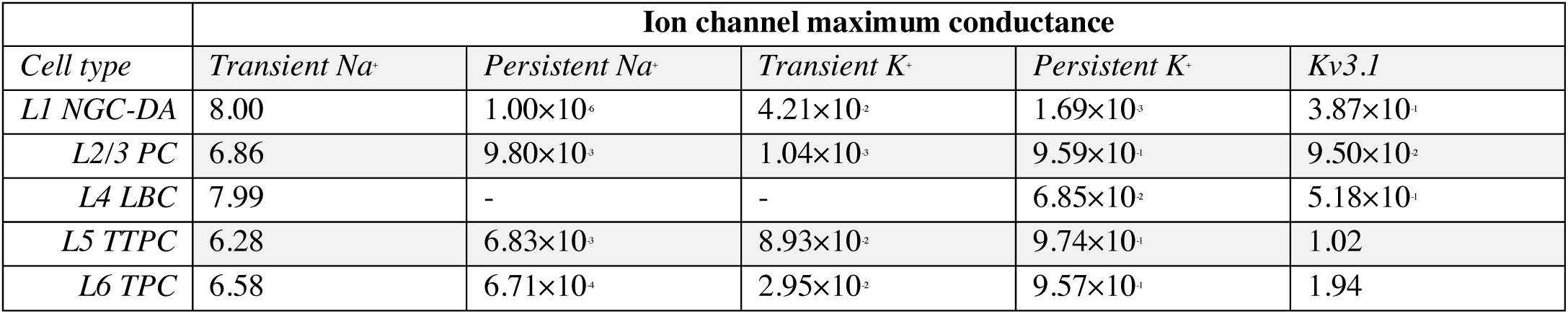
Maximal conductances in nodes and terminals (S/cm^2^). Conductances are the same across clones. All conductances are the same as in the original model axon initial segments, except Transient Na^+^, which was doubled.

Little is known about ion channel properties in cortical axon terminals, and even less is known about differences between cell types and cortical regions. Several studies found active sodium and potassium currents in pre-synaptic terminals [53–58]; therefore, terminations were assigned the biophysical properties of nodes of Ranvier [45,59]. The nodes and terminals had the same passive parameters as the initial segment, including a specific capacitance (*C*_m_) of 1 *μ*F/cm^2^ and cell-type-specific passive membrane resistance (*R*_m_), while the myelinated sections had an *R*_m_ of 1.125 MΩcm^2^ [60] and *C*_m_ of 0.02 *μ*F/cm^2^ [60–62].

#### 3.2.3 Adjusting Models to Reflect Properties of Adult Rat and Human Neurons

The original models were derived from juvenile rat cortical neurons; therefore, several geometric parameters were adjusted based on age and species related differences to represent adult rat and human neurons. For L5 thick-tufted pyramidal cells, total basal dendritic length increases from P14 to P60 by a factor of 1.17 and mean basal and apical dendritic diameters increase by factors of 1.13 and 1.25, respectively [25]. Somatic diameter increases by a factor of 1.32 from P14 to P42, after which changes were not statistically significant [26]. Axonal diameter is linearly related to somatic diameter [43,63], and axonal segment diameters were uniformly scaled with the somatic diameter.

Geometric scaling factors were derived by comparing the morphologies of previously published L2/3 pyramidal cells from adult human temporal cortex [23] to the Blue Brain L2/3 pyramidal cells. Median basal and apical compartment diameters were approximately 1.95 and 1.88 times larger, respectively, and somatic diameter was approximately 2.45 times larger in the human cells. The human neurons had more complex and numerous dendritic branching, that cannot be approximated by linear scaling [64], and we did not scale further the dendritic lengths. All the scaling factors are listed in Table 2.

**Table 2.** Scaling factors to transform juvenile rat cortical models to represent the geometry of adult rat and adult human cortical neurons.

| Morphological element | Scaling factor |  |
| --- | --- | --- |
|  | Adult, Rat | Adult, Human |
| Basal dendritic diameter | 1.13 | 1.95 |
| Basal dendritic length | 1.17 | 1.17 |
| Apical dendritic diameter | 1.25 | 1.88 |
| Somatic diameter | 1.32 | 2.45 |
| Axonal diameter | 1.32 | 2.45 |

After scaling, the adult rat models had somatic diameters from 10–20 *μ*m for all cells except the L5 pyramidal cells, which were 22–24 *μ*m in diameter (Figure 2b). Most axonal compartments were between 0.3–1 *μ*m in diameter, except for the axon hillocks and some proximal axonal branches that represented the upper diameter range (Figure 2c). Axon terminal compartments were mostly smaller than 1 *μ*m in diameter (Figure 2d), with a few exceptions that likely represented synaptic boutons or axon swellings (“blebs”) of compartments near the original slice boundaries. The relative diameter ranges between cell types was the same for the human cells, as the diameters were linearly scaled by the scaling factors in Table 2.

### 3.3 Electromagnetic Stimulation of Model Neurons

Using the quasistatic approximation [65,66], the electric field generated by neural stimulation can be separated into spatial and temporal components. The spatial component for each stimulation modality was applied to the compartmental models as extracellular potentials in NEURON using the extracellular mechanism [15,31,67] and scaled uniformly in time based on the time-dependent waveform. Analysis, visualization, and simulation control was conducted in MATLAB (R2017a, The Mathworks, Inc., Natick, MA, USA).

#### 3.3.1 Intracortical Microstimulation

ICMS was modeled as a point current source in a homogenous, isotropic medium with conductivity *σ* = 0.276 S/m [68]. For a microelectrode delivering current *I* centered at (*x*_0_, *y*_0_, *z*_0_), the extracellular potential *V*_*e*_ was calculated at the location of each compartment (*x*, *y*, *z*),

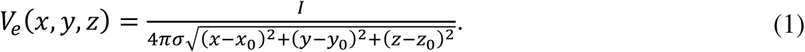

Thresholds were determined for cathodic pulses ranging in duration from *PW* = 50 *μ*s - 1 ms at electrode locations throughout a 3D grid encompassing each neuron’s full arborization in 100 *μ*m steps. Electrode locations within 30 *μ*m of the nearest neural compartment were considered too close for accurate calculation of extracellular potentials without considering the presence of the neural structures or the 3D structure of the microelectrode tip [69–72], and locations farther than 1 mm were outside of typical experimental ranges, so these points were removed from subsequent analysis. The chronaxie *τ*_*ch*_ and rheobase *I*_*rh*_ were obtained at each location by fitting equation (2) [73] to the thresholds using non-linear least squares regression in MATLAB,

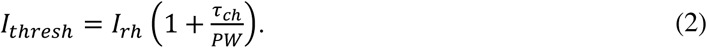

#### 3.3.2 Subthreshold and Suprathreshold Uniform E-field Stimulation

Transcranial electromagnetic stimulation modalities (TMS/tES) generate E-fields that have low spatial gradients at the scale of single neurons [74,75]. Therefore, we applied a uniform E-field to the model neurons, and the field was parameterized by only an amplitude and the angle relative to each neuron’s somatodendritic axis. The extracellular potential at each compartment (*x*, *y*, *z*) in a uniform E-field with direction given by polar angle *θ* and azimuthal angle *ϕ*, in spherical coordinates, and the origin (soma) set to ground, was calculated using equation (3):

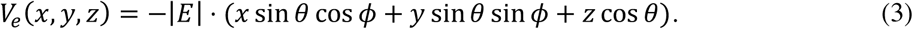

After calculating the E-field distribution, amplitudes were scaled over time by various current waveforms. To simulate subthreshold polarization by transcranial direct current stimulation (tDCS), 100 ms rectangular pulses were applied at an amplitude of 1 V/m [6]. For suprathreshold simulations, threshold amplitudes were obtained for rectangular pulses, ranging from 10 *μ*s to 100 ms in duration.

### 3.4 Simulation Parameters

The neural models were discretized with isopotential compartments no longer than 20 *μ*m and solved using the backward Euler technique with a time step of 5 *μ*s or 1 *μ*s for threshold simulations with pulse durations of 50 *μ*s or less. Ion channel kinetics were scaled to 37°C using a *Q*_10_ of 2.3. After myelinating the axons, the number of compartments ranged from 609 for one of the L2/3 PCs to 4,048 for one of the densely arborized L4 LBCs. Thresholds for AP generation were determined using a binary search algorithm with 0.5 V/m (and 0.5 *μ*A) accuracy; activation was defined as the membrane potential crossing 0 mV (with positive slope) in at least 3 compartments within the neuron. Before stimulation was applied, the membrane potential of each compartment was allowed to equilibrate to steady state.

## 4. Results

First, we simulated the responses of the original and modified model neurons to dc current injection in the soma (4.1). Subsequently, we simulated three forms of cortical stimulation and compared the responses of the modified models to published experimental data for intracortical microstimulation of L5 pyramidal cells (4.2), subthreshold polarization in a uniform dc E-field (4.3), and suprathreshold stimulation with a uniform pulsed E-field (4.4). Finally, we quantified differences in stimulation responses between the adult rat and human models (4.5).

### 4.1 Firing Behavior of Modified Model Neurons

The parameters of the original models were optimized with only the axon initial segment included. To quantify the changes in firing properties resulting from to our modifications, particularly the inclusion of myelinated axonal arbors, we compared frequency-current (F-I) curves of the baseline models and the myelinated, scaled versions (Figure 3). Markram et al. classified each neuron into electrical types based on their characteristic response to step current inputs, e.g., continuous adapting or burst accommodating [29]. The characteristic response types of the model neurons did not change after modifications (example traces in Figure 3); however, the F-I curves of the modified models were generally less steep and shifted rightward, which is consistent with the decrease in somatic input resistance observed between immature and mature cortical neurons [27]. Some of the model neurons exhibited shortened AP heights and depolarization block at higher current amplitudes. In these model neurons, increasing the temperature from 34° to 37° C caused depolarization block to occur at lower current amplitudes, due to the faster Na^+^ inactivation kinetics.

**Figure 3.**
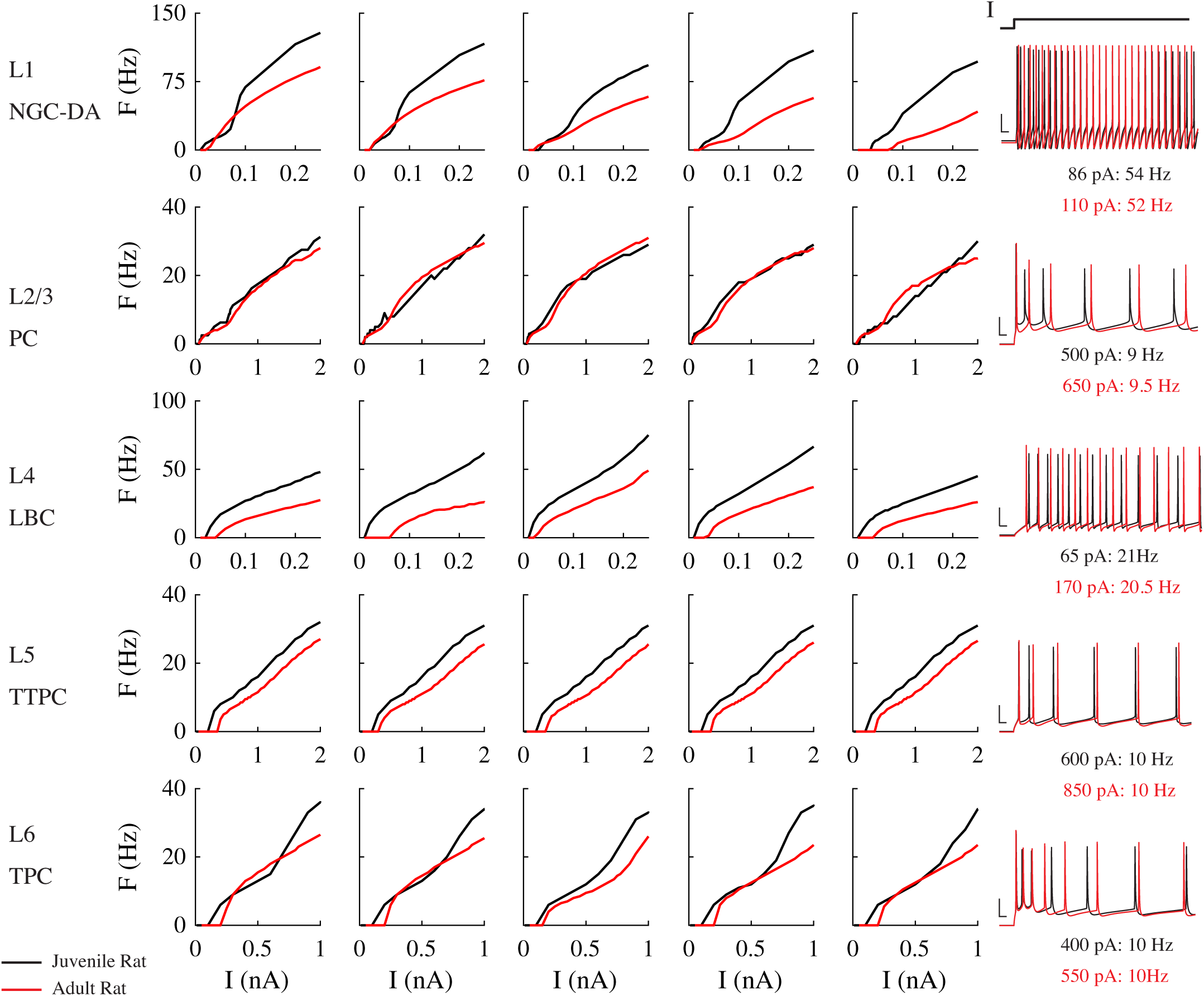
Firing behavior in modified and original cortical model neurons. Frequency-current (F-I) curves for original, juvenile rat Blue Brain models (black) and modified, adult rat models (red). Original models were unscaled and included only 60 *μ*m axon initial segment (AIS). Modified models were scaled up in size to approximate adult rat cortical neurons and included myelinated axonal arbors, as described in Section 3.2. Temperature set to 34° C for modified models to match [29]. Each row includes the F-I curves for the 5 virtual clones and membrane potential recordings in the original and modified models at current amplitudes producing similar firing rates.

We also compared the F-I curves of model L2/3 PCs to *in vitro* recordings of L2/3 PCs from human temporal cortex [64] (Figure 4). Scaling the models to human size further increased the threshold current to initiate at least one action potential (AP) and decreased the slope relative to the adult rat models, but did not increase the threshold as high as recorded experimentally in human neurons. Beyond the reference current producing firing at 10 Hz, the F-I curves of the human model neurons were closer to the mean of the experimental recordings than the adult rat model neurons.

**Figure 4.**
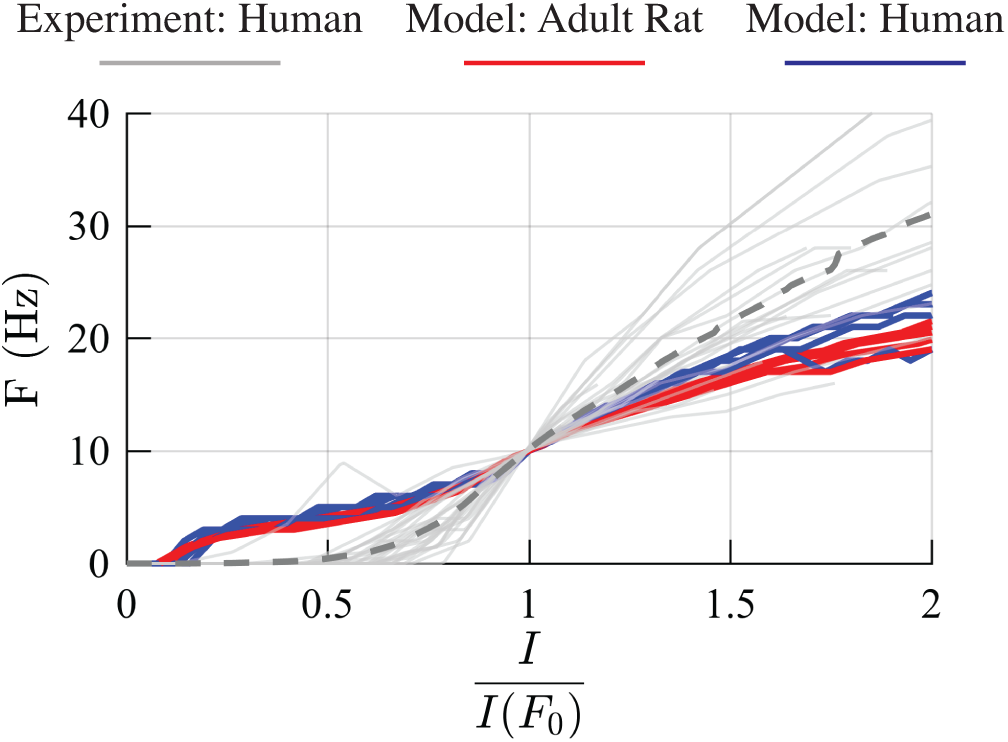
Firing behavior of human-scaled L2/3 PC model neurons. F-I curves for the five adult rat (red) and human (blue) L2/3 PCs models, as well F-I curves for 25 human L2/3 PCs recorded *in vitro* [63]. Mean F-I curve for experimentally recorded neurons shown in dashed gray line. Current axis normalized to current amplitude producing reference firing rate *F*_0_ = 10 Hz.

### 4.2 Intracortical Microstimulation of Model L5 Pyramidal Neurons

We calculated threshold current amplitudes for 50 *μ*s to 1 ms duration cathodic stimuli applied across a 3D grid of electrode locations surrounding each of the five L5 pyramidal cell models. APs were always initiated in axonal compartments—either at a node of Ranvier or an axon terminal—and never in the soma or dendritic arbor; Figure 5a illustrates the locations of AP initiation for an example cell (L5 PC 1). When the electrode was placed immediately adjacent to the apical dendrites, the closest dendritic compartments were passively charged past the defined threshold, but despite the presence of active channels, they did not initiate propagating APs. Within the axonal arbor, ICMS predominantly activated axon terminals compared to nodes of Ranvier along axon branches (Figure 5b); the latter instances occurred when the electrode was positioned close to a node of Ranvier with no nearby terminals.

**Figure 5.**
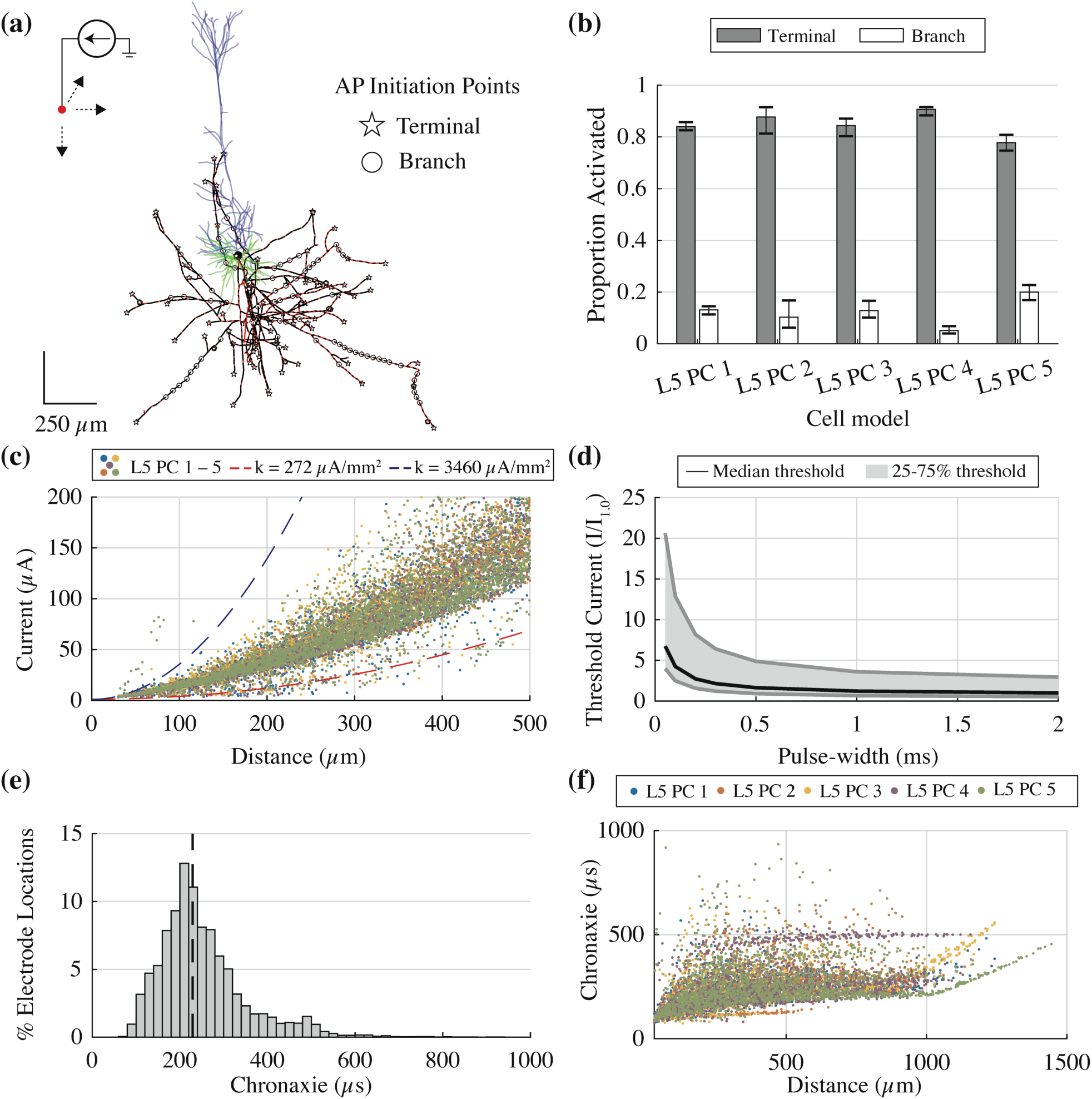
Model L5 pyramidal cells reproduced experimental responses to ICMS. **(a)** AP initiation sites for 0.2 ms pulse, all electrode positions, shown for L5 PC 1. **(b)** Mean proportion of AP initiation points at either axon terminal or branch across pulse durations (50 *μ*s to 1 ms). Error bars are minimum and maximum. **(c)** Threshold current-distance plot for all five L5 pyramidal cells with 0.2 ms cathodic pulse. Each point represents the threshold for a different electrode location within the 3D grid and the distance from that electrode to the point of action potential initiation. The current-distance curves are plotted for current-distance constants of cat pyramidal tract cells [75]. **(d)** Strength-duration curves for median (black line) and 25 - 75 percentile thresholds (gray shaded region). Median chronaxie ± standard deviation was 229 ±101 *μ*s. (**e)** Histogram of chronaxies at all electrode locations for 5 L5 PC models. Black dashed line indicates median chronaxie value (229 *μ*s). **(f)** Chronaxies plotted against distance from electrode location to AP initiation site for 5 L5 PC models.

The threshold current for electrical microstimulation to activate a neuron is proportional to the square of the distance between the stimulating electrode and the neuron [76]. This current-distance relationship can be expressed as *I*(*r*) = *kr*^2^ + *I*_0_, where *I* is the threshold current, *r* is the distance, *I*_0_ is the minimum threshold, and *k* is the current-distance constant, in *μ*A/mm^2^. We determined the distance between each electrode location and the compartment in which the AP was initiated for the 0.2 ms cathodic pulse in all five L5 PCs (Figure 5c); the threshold points fell largely within the experimental range for L5 pyramidal tract neurons in cat motor cortex (*k* = 272 - 3460 *μ*A/mm^2^) [77]. Fitting the current-distance equation to the threshold points for all five cells using regression yielded a *k* value of 483 *μ*A/mm^2^ (95% confidence interval 480 - 485 *μ*A/mm^2^). The threshold values that fell below the experimental threshold range occurred at electrode locations where activation occurred through mechanisms other than direct cathodal activation, e.g., anode break effects at the virtual anodes [75,78,79], and the points above the experimental threshold range resulted from activation of unmyelinated branches.

Figure 5d depicts the normalized strength-duration curve for the median and 25^th^–75^th^ percentile thresholds across all electrode positions. We fit the Weiss equation (2) to the thresholds at each electrode location and obtained a median chronaxie (± standard deviation) of 229 (± 101) *μ*s (Figure 5e). These data fall within the 100 - 20 *μ*s chronaxie range measured for single L5 pyramidal tract neurons in cats [77,80], and the 100 - 300 *μ*s range obtained using behavioral methods in non-human primates [81]. One of the factors underlying the heterogeneous distribution of chronaxies was the distance from electrode location to AP initiation site: electrode locations that activated more distant compartments exhibited longer chronaxies (Figure 5f), especially beyond 1 mm, and distance and chronaxie had a Pearson correlation coefficient of 0.38.

### 4.3 Subthreshold Stimulation with Uniform dc E-field

We applied downward, uniform E-fields (*θ* = 180°) to each model neuron with 100 ms pulses and calculated the change in transmembrane potential in each compartment. This configuration matches the experimental setup of Radman et al. where brain slices from rat motor cortex were stimulated with uniform E-fields with the anode on the pial surface and the cathode opposite [6]. The model neurons exhibited a biphasic distribution of membrane polarization, with hyperpolarization near the anode and depolarization near the cathode (Figure 6a), as observed experimentally [82,83]. We computed *polarization lengths* for each model neuron by dividing the change in transmembrane potential at steady state by the magnitude of the E-field, with units of mV/(V/m). The polarity of the somatic polarization length depended on the soma’s relative position between the two ends of the cell, and the magnitude depended on the diameter, length, and orientation of the dendritic and axonal branches. L5 pyramidal cells had the largest somatic polarization lengths followed by L2/3 pyramidal cells and interneurons, in agreement with *in vitro* measurements (Figure 6b). The range of somatic polarization magnitudes in the model neurons was smaller than in the experiments, likely due to the reduced number and variability of the cell morphologies tested in our simulations.

**Figure 6.**
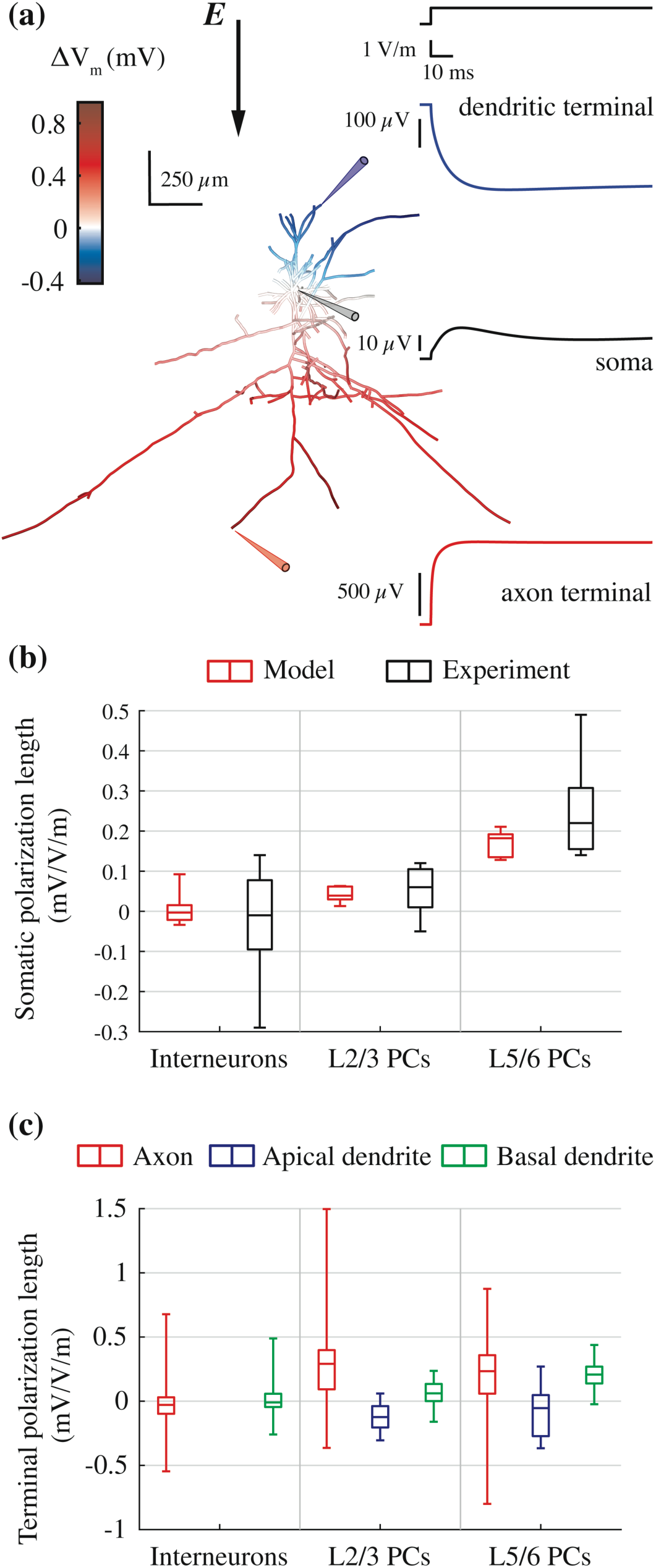
Model neurons reproduced experimental responses to subthreshold uniform E-field stimulation. **(a)** Color plot of membrane polarization in example L2/3 PC at end of 100 ms uniform E-field directed downward (left) and recordings from a dendritic terminal, the soma, and an axonal terminal indicated by the colored probes (right). **(b)** Box-plots of somatic polarization lengths for downward E-field (*θ* = 180°) of model (red) and experimental data (black) from [6]. Experimental data from neurons with dendritic arbors cut during brain slice preparation were excluded. **(c)** Box-plots of terminal polarization lengths for axons, apical dendrites, and basal dendrites.

Within each model neuron, axon terminals experienced the largest polarization, with maximal polarization at longer, distal branches parallel to the E-field (Figure 6c). The maximal polarization length for L5/6 PC axon terminals was 0.88 mV/(V/m), which agrees well with the maximum polarization of 0.8 mV/(V/m) for axon terminals (blebs) by uniform E-fields in mouse cortical slices [4]. The median axon terminal polarization of L2/3 PCs (0.29 mV/(V/m)) and L5 PCs (0.23 mV/(V/m)) was similar, while the maximal L2/3 PC terminal polarization reached 1.50 mV/(V/m) (Figure 6c). This is explained by the longer vertical, intracortical axon branches of the L2/3 PCs (Figure 1) as both theoretical derivations and experimental measurements show terminal polarization increases with the length of the final axonal branch [4,9]. Despite negligible somatic polarization, model interneurons experienced significant polarization at their axon terminals. Finally, apical and basal dendritic terminals of all cell types experienced larger magnitude, faster polarization responses than the cell bodies, consistent with *in vitro* measurements [82].

### 4.4 Suprathreshold Stimulation with Uniform Pulsed E-field

We simulated direct activation by pulsed uniform E-fields to study the effects of morphological factors on model neuron responses to suprathreshold transcranial stimulation modalities. Thresholds for 10 *μ*s – 100 ms duration rectangular pulses were calculated for E-field directions spanning the polar (*θ*) and azimuthal (*ϕ*) directions in steps of 15° and 10°, respectively. Across cell types, the sites of maximal depolarization and AP initiation for the minimum-threshold orientation were at axonal terminations aligned with the E-field. Changing the E-field direction (or, equivalently, the cell orientation) changed the threshold and shifted the activation site (Figure 7a). APs were not initiated at the soma or axon initial segment for any orientation or pulse duration tested, and APs initiated at a terminal propagated antidromically back to the cell body and re-orthodromically throughout the axonal arbor (Figure 7b).

**Figure 7.**
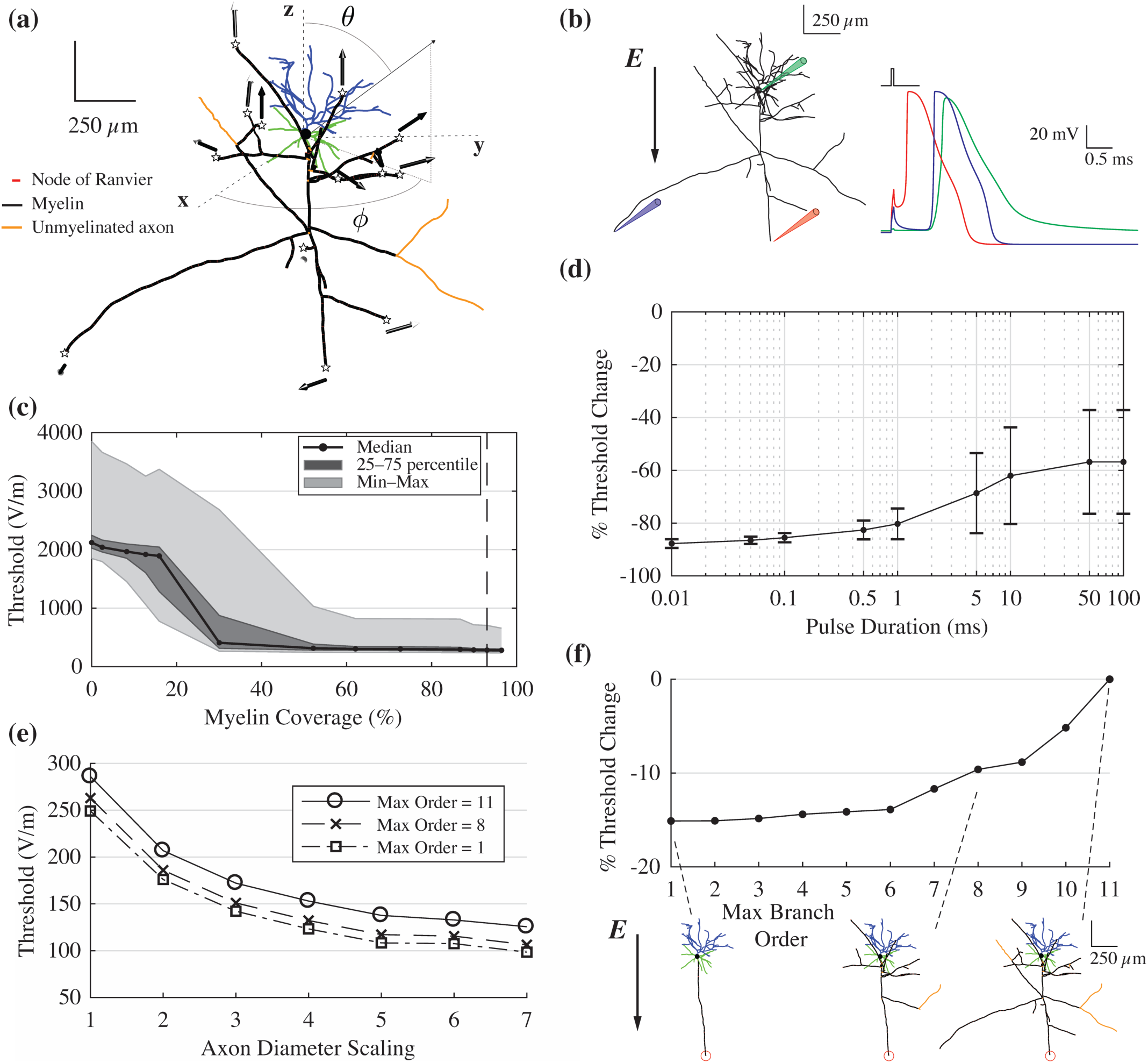
Threshold dependence on morphology with uniform E-field stimulation. **(a)** Unique AP initiation sites (stars) overlaid on morphology of example L2/3 PC for full range of uniform E-field directions using 50 *μ*s rectangular pulse. Arrows correspond to E-field direction for lowest-threshold AP initiation, with arrowhead color indicating the vector points out of the page (black) or into the page (white). **(b)** Membrane potential recordings from example L2/3 PC model neuron at AP threshold for 50 *μ*s rectangular pulse with E-field direction indicated by black arrow. Recordings made from directly activated axon terminal, indirectly activated axon terminal and soma. **(c)** Threshold of L2/3 PC with 50 *μ*s rectangular pulse across E-field directions as percent myelin coverage is varied **(d)** Percent change in threshold after adding myelin to axon of L2/3 PC (mean ± SD across E-field directions) for rectangular pulses of increasing duration. **(e)** Threshold E-field amplitude as diameter of axonal arbor uniformly scaled up for L2/3 PC with the three levels of arborization shown in (f): maximum axon order of 1 (square), 8 (cross), and 13 (open circle). Same E-field direction as in (b). **(f)** (top) Percent reduction in threshold for terminal activation (indicated by red circle) as axon collaterals were successively removed by decreasing branch order. (bottom) Neuron morphologies for maximum axon branch orders of 1, 8, and 11. Same E-field direction as in (b),

We analyzed the effects of myelination, axon diameter, and the degree of arborization to understand better the morphological factors that determine the thresholds for E-field pulses. We varied the degree of myelin coverage by adjusting the minimum axon diameter and maximum branch order to myelinate (Figure S2) and found that myelination dramatically reduced thresholds for all E-field directions (Figure 7c). Once approximately 60% of the axonal arbor was myelinated, additional myelination had almost no effect on thresholds across E-field directions. The effect of myelin on threshold was pulse-duration-dependent: myelination reduced thresholds more for shorter pulse durations; monophasic TMS pulses are similar to rectangular pulses in the 50–100 *μ*s range, corresponding to a mean threshold reduction of ∼86% (Figure 7d). Uniformly increasing the diameter of the entire axonal arbor also reduced threshold with an approximately inverse-square-root relationship (Figure 7e). Finally, removing higher-order collaterals (pruning) while retaining the main axonal branch reduced thresholds by as much as 10%, indicating that more local branching near an activated terminal increases the threshold (Figure 7f). These results (Figure 7) are from one of the L2/3 pyramidal cell models, but the trends were qualitatively identical across model neurons.

The complex dependence of threshold and AP initiation site on axonal morphology was a direct result of the realistic, branched arborization. We compared the effect of E-field direction on activation threshold in one of the L2/3 PC models to a commonly used L3 PC model with reconstructed dendritic morphology and an artificial, straight axon [61] (Figure 8a). The threshold of the realistic model was much less sensitive to E-field direction than the straight axon model, due to the latter having only a single axon terminal capable of activation (Figure 8b). The threshold of the straight axon model was maximal when the E-field was perpendicular to the somatodendritic axis with a mean of 21.8 (normalized to minimum), averaged across azimuthal rotations. In contrast, the realistic axon model was much less sensitive to E-field rotations in the polar direction, while azimuthal rotations produced wider variations in threshold relative to the mean. The threshold of the straight axon model dropped sharply when the E-field was oriented downward, aligned with the axon; to a lesser degree, there was also a slight preference for downward oriented fields in the realistic model.

**Figure 8.**
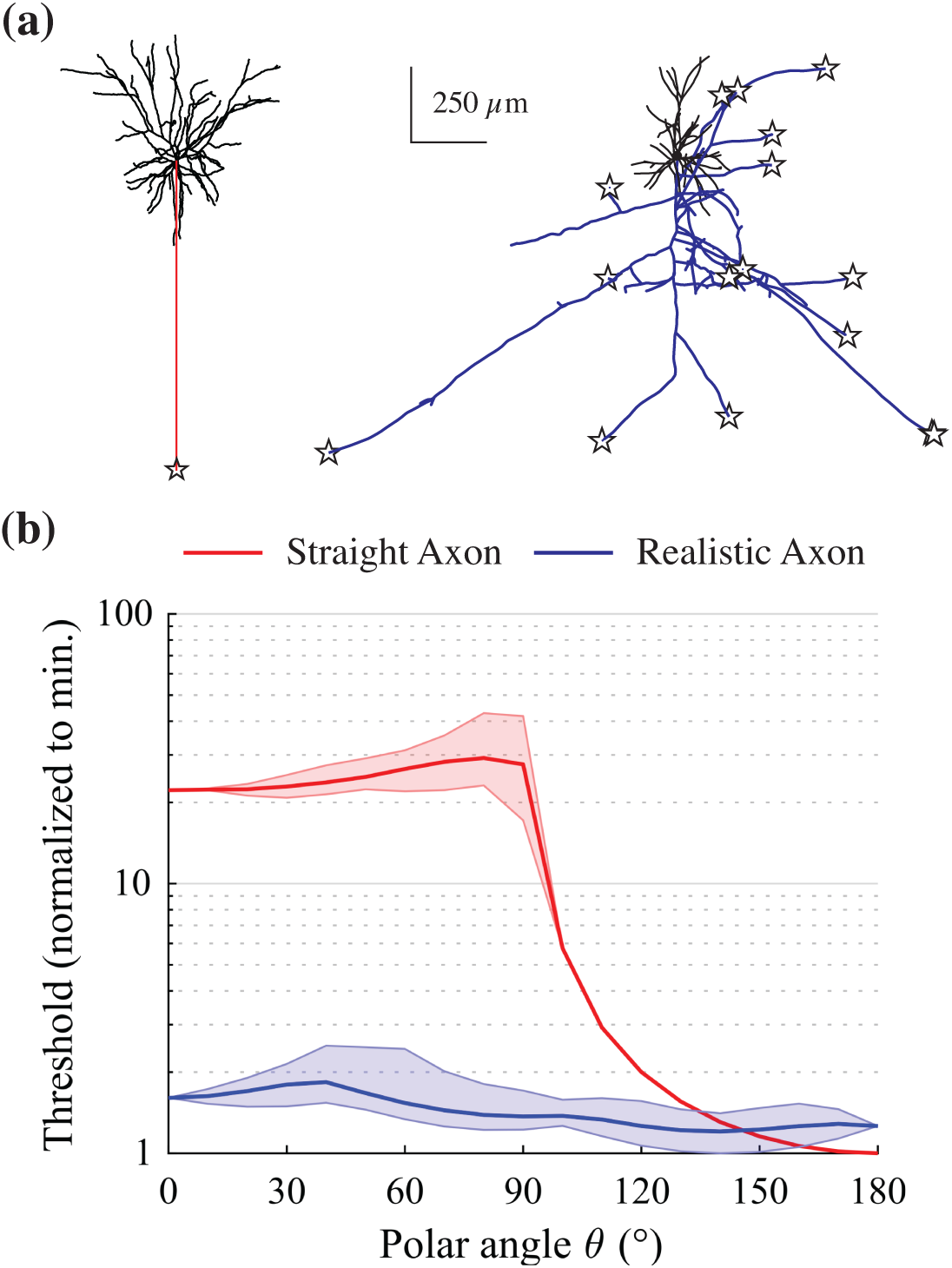
Threshold dependence on E-field direction with branched vs simplified axon morphology. **(a)** Morphologies of L2/3 PC with branched axon (left) and with simplified, linear axon (right), with unique AP initiation sites overlaid as stars. Model with simplified axon drawn from [60] **(b)** Normalized threshold as polar angle *θ* of E-field direction varied. Dark middle line is mean threshold, and shaded region is range, across all azimuthal angles *ϕ*.

Radman et al. also applied uniform E-fields to rat cortical slices with 100 ms pulses at suprathreshold intensities, and found that the spiking responses were always produced following excitatory post-synaptic potentials (EPSPs). The minimum E-field intensity at which EPSPs were observed was between 12 and 104 V/m, which provides an indirect estimate of the threshold range of pre-synaptic axonal elements in the slice. The model threshold intensities for activation of axon terminals largely fell within this experimental range, with some E-field directions producing APs at intensities slightly below 12 V/m for the L2/3 and L5 PCs (Figure S3).

### 4.5 Comparison of Human and Rat Model Neuron Responses to Stimulation

While the mature rat- and human-scaled model neurons had qualitatively similar responses to ICMS and uniform E-field stimulation, there were quantitative differences in the measures of excitability and polarization. In response to ICMS, the human L5 PC models had ∼100 *μ*A/mm^2^ lower current-distance constants and 30 *μ*s shorter median chronaxies than the adult rat L5 PCs, indicating reduced thresholds for activation and faster membrane responses (Figure 9a). The range of somatic polarization lengths for uniform E-field stimulation with 100 ms pulses was increased on the order of a few *μ*V/(V/m), and the range of axonal and dendritic terminal polarization lengths increased 20–171 and 2–36 *μ*V/(V/m), respectively (Figure 9b). Finally, we compared the activation thresholds between cell types for a 50 *μ*s rectangular pulse applied at the same range of E-field directions as in Figure 7. As with ICMS, the human model neurons had consistently lower thresholds than the rat models, retaining the same threshold trends between cell types within both sets of model neurons (Figure 9c).

**Figure 9.**
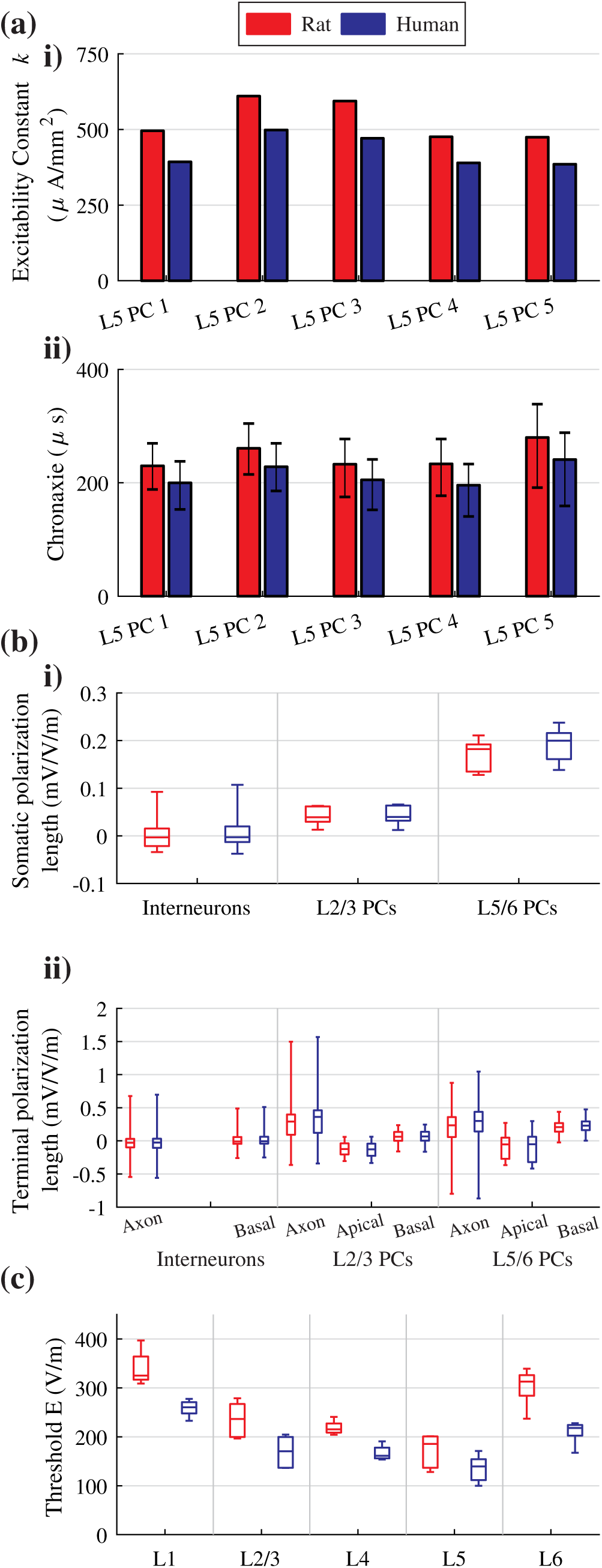
Human and rat model response to different stimulation modalities. **(a)** ICMS results for rat (red) and human (blue) L5 PC models. **i)** Current-distance constant *k* fit for each L5 PC model using linear regression. Electrode locations within 30 *μ*m of closest neural compartment and outside of 1 mm excluded. **ii)** Median chronaxie for each L5 PC model. Error bars correspond to standard deviation. **(b)** Box-plots of polarization lengths for downward E-field (*θ* = 180°), rat (red) and human (blue) model neurons. **i)** Somatic polarization lengths. **ii**) Terminal polarization lengths of axons, apical dendrites, and basal dendrites. **(c)** Box-plots of threshold amplitudes for 50 *μ*s rectangular pulse across all E-field directions for rat (red) and human (blue) model neurons.

## 5. Discussion

We implemented a library of morphologically-realistic compartmental models of cortical neurons modified to reflect the morphology of adult rat and human cortical neurons and validated the excitation properties of the models across several modalities of cortical stimulation. The original models were drawn from the Blue Brain network model of juvenile rat somatosensory cortex and were adapted to approximate the geometrical differences of mature cortical cells in rats and humans, including extensive myelinated axonal arbors. We characterized the model responses to ICMS as well as sub- and supra-threshold regimes of uniform E-field stimulation, revealing agreement with multiple sources of experimental data. The models reproduced the current-distance and strength-duration relationships measured *in vivo* with ICMS, as well as somatic and axonal polarization in uniform E-fields measured *in vitro* in brain slices. Additionally, these models reinforced previous evidence of the preferential polarization and activation of axon terminals by both intracortical and transcranial stimulation. Finally, the results suggest incorporating realistic axonal geometries is critical to predicting accurately variations in polarization and activation thresholds with spatial and temporal parameters of stimulation, as well as variations between and within cell types.

### 5.1 Activation of Axon Terminals

Axon terminals were the most likely neural element to be activated by ICMS, and uniform E-field stimulation produced maximal polarization and initiated APs at axon terminals aligned to the E-field. These results can be explained by cable theory: extracellular E-field stimulation produces depolarization proportional to the parallel component of the E-field gradient, known as the activating function [12,84]. An externally applied E-field therefore generates sources or sinks of transmembrane current in neuronal compartments based on the interaction between the neural geometry and the E-field distribution [85]. In uniform E-fields, depolarization is predicted to occur only at sites of geometrical discontinuities—including terminations, bifurcations, and bends— and is driven by the E-field magnitude, rather than the E-field gradient [75,86]. Even in non-uniform E-fields, these discontinuities are potential sites of strong polarization when appropriately situated. Of these discontinuities, axon terminals should experience the greatest polarization, because unlike bends and branch points, shunting via axial paths of current flow is minimized at terminals, where only a single path exists for current to flow away from the polarized membrane [87,88].

Recordings from axon blebs provide the first direct, experimental measurements of axon terminal polarization by uniform dc E-field stimulation [4]. The range of axon terminal polarization in our L5 PC models agreed with these measurements to within 10%, while the maximal L2/3 PC terminal depolarization exceeded that of the L5 PCs and the experimental estimates. This can be explained by theoretical derivations showing that terminal polarization is proportional to diameter and final branch length [9] and differences in the axon branch orientations relative to the E-field: the L2/3 PCs possessed longer vertical and oblique axon branches, extending downward to deeper cortical layers, which were maximally depolarized for the applied, downward E-field direction, while the L5/6 PC axons had longer branches extending to superficial layers, which experienced the greatest hyperpolarization. The relative axonal diameters and lengths were responsible for the polarization trend between L2/3 and L5/6 PCs, as well as the smaller range of terminal polarizations in the finer, more locally arborized interneuron axons. The slight overestimate of maximal L5 PC axon terminal polarization was likely due to the larger diameter axon terminals of the adult rat models relative to the mouse neurons used in the experiments. The peak E-field in the cortex during 2 mA tDCS in humans was measured to be approximately 0.8 V/m *in vivo* [89] and 0.33 V/m *post mortem* [90]. Taken together with our modeling results in the human neuron models, this suggests tDCS may polarize some cortical axon terminals by 0.5–1.3 mV. Further support for these findings will require more detailed modeling of the E-field distribution in the brain (see Section 5.2).

Consistent with polarization by subthreshold fields, axon terminals were the most excitable neural elements for suprathreshold stimulation, in agreement with previous modeling studies of finite axons [75,86], cortical neurons with idealized morphologies [7], and cortical neurons with realistic morphologies [8]. The latter study by Wu et al. used methods similar to the present study to simulate models of L2/3 and L5 PCs with myelinated axon collaterals. Like our results, they demonstrated that axon terminals are preferentially activated by pulsed stimulation, that the threshold was inversely related to the axon diameter, and that the threshold was directly related to the degree of branching. However, some of their models were activated at the soma for certain E-field directions and long pulse durations, which we did not observe. This may be related to differences in AP generation caused by the uniform ion channel conductances used across cell morphologies or the ion channel models drawn from an older study with different morphologies [91]. In addition, Wu et al. flattened the cellular morphologies into 2D and used a simpler approach to myelination that did not allow for nodes of Ranvier between branch point; our results demonstrated that these geometrical changes can have profound effects on threshold.

Multiple lines of experimental evidence implicate axon terminals as particularly sensitive to electrical stimulation. Radman et al. found *in vitro* that nearly all APs evoked by uniform dc E-fields were preceded, at lower field strengths, by excitatory post-synaptic potentials due to activation of afferent inputs [6]. Blocking glutamatergic synaptic transmission abolished the EPSPs and cells no longer fired APs in response to stimulation. Similarly, Nowak and Bullier not only demonstrated that microelectrode stimulation activates axons, rather than cell bodies, but also provided evidence that direct activation of local cells occurs predominantly at axonal branches [33,92]. Hussin et al. reported similar results in rat motor cortex, both in brain slices and *in vivo*: blocking synaptic transmission caused a significant reduction in the spiking response and an increase in the current threshold to evoke movements [93]. Lastly, several *in vivo* studies in non-human primates and cats found that microelectrode stimulation activated afferent inputs at lower thresholds than local cells [80,94–96]. These data are explained by the low threshold of axon terminals and branch points to electrical stimulation observed in our models, which leads to predominantly indirect activation via afferent inputs.

### 5.2 Effect of Axon Morphology on Direct, Cortical Response to Stimulation

Incorporating cell-type-specific axonal arbors determined how activation thresholds varied with stimulation parameters. Previously, the only studies to simulate ICMS of models with branched, axonal arbors used artificially generated or flattened axon morphologies with limited spatial extent [20,21]. The axons in our models included full representations of local axonal arborization, which is especially dense for interneurons, as well as mid-range, horizontal projections of pyramidal cells (1 - 2 mm), but did not include long-range projections (> 2 mm) [42,97,98]. The variability in initiation site, current-distance constant (*k*), and chronaxie with electrode locations relative to the axonal arbor suggests that the population input-output curves at different electrode depths computed in prior studies [20,21] would be altered with more realistic axon morphologies. Furthermore, some studies suggested that ICMS directly activates local cells at their “initial segment and nodes of Ranvier” [76], but our biophysically-realistic models suggest that local cells are activated predominantly at their axon terminals or nodes of Ranvier beyond the initial segment; this finding is supported experimentally by the wide distribution of low threshold points for ICMS of single pyramidal tract neurons [80].

Our results also indicate that myelination significantly reduces thresholds, with larger reductions for briefer pulses. This can be explained by taking into account the influence of myelin on the effective length constant [67,85] as well as the frequency-dependent spatial decay of membrane polarization along the cable [99]. The “effective” length constant increases with inclusion of myelin and decreases as a function of frequency. Therefore, despite uncertainties about the properties of intracortical myelination, discussed in Section 5.3, we expect it to have a significant effect on thresholds for stimulation modalities with short pulse durations (e.g. TMS).

Additionally, previous studies coupled compartmental neuron models to finite element method (FEM) head models to predict the direct cortical response to TMS. However, these neuron models lacked realistic axon morphologies with intracortical branching, resulting in exceedingly high thresholds [15] or unlikely sites of AP initiation, e.g., axon initial segment [18]. Excluding axon collaterals produces significant differences in the orientation (directional) dependence of threshold (Figure 8), and this is a critical factor for calculating the neural response to changes in the spatial parameters of stimulation, e.g., TMS coil orientation. The inclusion of realistic intracortical axonal arbors for a range of cell types makes the present models well-suited for coupling to 3D FEM models of the electric field distribution. These coupled models can be used to quantify polarization and activation of neurons throughout the cortical layers for a variety of brain stimulation modalities.

### 5.3 Model Limitations

Our models build on previous work by adapting multi-compartmental neuron models with realistic axonal arbors. The resulting model neurons were validated against a range of electrophysiological recordings, but have several limitations that should be noted.

One difficulty in estimating *in vivo* activation thresholds and measures of excitability in models of single neurons arises from differences in the state and electrophysiological properties of *in vivo* and slice preparations, the latter of which are the source of most modeling data. We modeled quiescent neurons, which may overestimate thresholds for neurons receiving ongoing synaptic inputs, and some evidence suggests there are further differences between *in vitro* and *in vivo* neural properties, caused by other factors, such as neuromodulators or extracellular ion accumulation [100].

The Blue Brain models were derived from juvenile rats, and cortical axons undergo cell-type-specific pruning and growth as they mature [25]. In agreement with previous modeling work, we found that higher degrees of branching increased the thresholds of axon terminals, and determining objective methods of “aging” axonal morphology or using fully intact reconstructions from mature animals may be important for obtaining accurate activation thresholds. Additionally, aging is also associated with electrophysiological changes, including faster AP dynamics [27] and burst-firing behavior in L5 TTPCs, likely mediated by increased dendritic ion channel densities [101,102]. Human pyramidal cells also have more complex dendritic trees, faster AP onsets, and are capable of firing at higher frequencies than rodent and even non-human primate pyramidal cells [24,103]. Human cortical neurons may even have a lower specific membrane capacitance than that of rodents and non-human primates [23], which will also affect their response to stimulation. These species-related differences may have contributed to the differences between the F-I curves of our human-scaled models and *in vitro* data from human cells (Figure 4).

The model neurons were drawn from rodent somatosensory cortex and did not include large pyramidal tract neuron or Betz cell morphologies found in motor cortex, and these neurons mediate the majority of corticospinal activity evoked by TMS and suprathreshold tES [104]. These cells may warrant more specific model development to simulate motor cortex stimulation due to their unique morphological and electrophysiological properties [105–108]. Although we accounted for and demonstrated the importance of age- and species-related differences in myelination, diameter, and basal dendritic length, addressing the additional factors identified above may be important to improve further the fidelity of models of human and rat neurons at different stages of development.

There are few data on the properties of intracortical axons, and we used the ion channel models and optimized conductances from the axon initial segments in the original models to represent the nodes in the entirety of the axon. These include a fast sodium channel model developed to explain the lower AP threshold in the axon observed experimentally by incorporating a shift in the activation curves between the initial segment and soma [109]. However, this model features slower kinetics and a lower conductance ratio between the axon initial segment and the soma than models based on more recent studies of axonal excitability [110–113]. While unlikely to alter the main conclusions of this study, more accurate axonal ion channel models may produce lower thresholds for electrical stimulation, particularly with short pulse durations.

As discussed earlier (Section 5.1), multiple lines of evidence support that intracortical axon terminals are directly modulated by nearly all stimulation paradigms, but we still lack clear knowledge of the geometrical and biophysical properties of intracortical axon terminals and nodes of Ranvier. Myelination was limited by a minimum axon diameter and branch length based on the literature, but the extent of myelination may have been overestimated, particularly near pre-synaptic terminals. The precise distribution of myelination on intracortical axons remains uncertain: sub-populations of pyramidal cells may have distinct patterns of myelination [36], and the structure and function of myelination near pre-synaptic terminals is still being determined [54]. We also did not incorporate terminal or *en passant* synaptic boutons in the axonal morphologies, which may increase the activation thresholds of axon terminals, as the increased surface area would reduce the input resistance [71]. Further experimental study of the electrophysiological and morphological properties of cortical axons is critical for the advancement of models and, ultimately, our understanding of cortical stimulation.

## 6. Conclusion

We generated biophysically-realistic models of adult rat and human cortical neurons that incorporate three-dimensional, morphologically-reconstructed axonal arbors, cell-type-specific electrophysiology, and species-specific adjustments. These models represent significant improvements over previous models used to study electromagnetic brain stimulation. The realistic morphologies make these models well-suited for simulations of cortical stimulation that incorporate detailed computations of the spatial distribution of the E-field. Our results demonstrate the sensitivity of axon terminals to both intracortical and transcranial methods of cortical stimulation, and this finding has important implications for the mechanisms of cortical stimulation. Finally, we presented results from 25 models (5 cell types) extracted from the library of publicly-available Blue Brain neuron models, and our modeling framework can be applied to any model from that library of 207 unique cell types.

## Acknowledgements

This work was supported by NIH grants R01 NS088674, R01 NS095251, and R25 GM103765 (Duke BioCoRE), and NSF Graduate Research Fellowship DGF 1106401. We thank Dr. Boshuo Wang, Dr. Andreas Neef, Dr. Stefan Goetz, Dr. Marc Sommer, and Karthik Kamaravelu for their technical assistance and helpful discussions. We also would like to thank the Duke Compute Cluster team for computational support.

## Supplementary Figures

**Figure S1.**
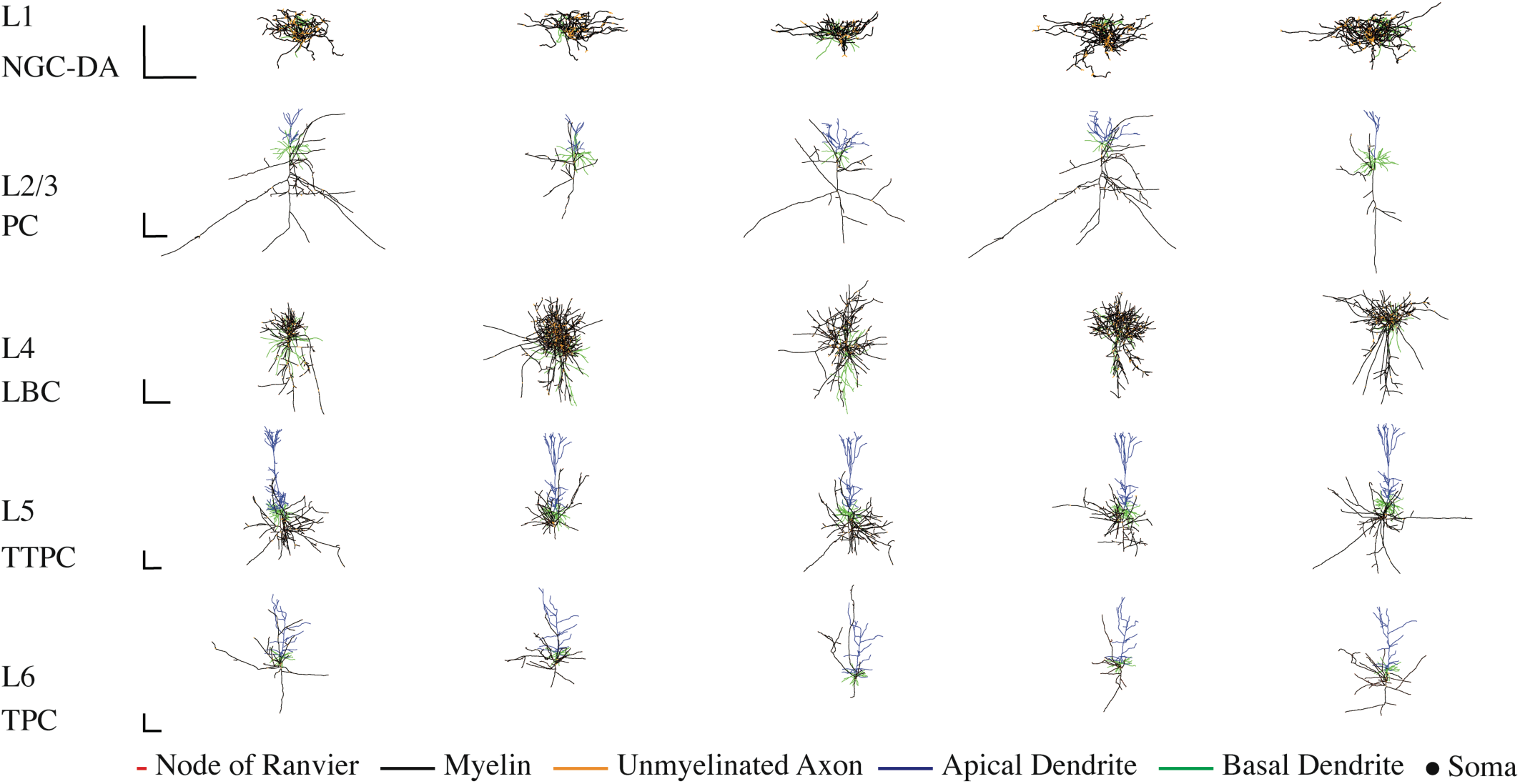
Morphology of model human cortical neurons. 3D reconstructions of morphological cell types used in this study. Each row contains the 5 clones of each cell type drawn from the layer indicated on the left. Adult human versions shown here, with morphologies colored to indicate nodes of Ranvier (red), myelin (black), and unmyelinated axonal sections (yellow), as well as apical dendrites (blue) and basal dendrites (green). Scale bars = 250 *μ*m.

**Figure S2.**
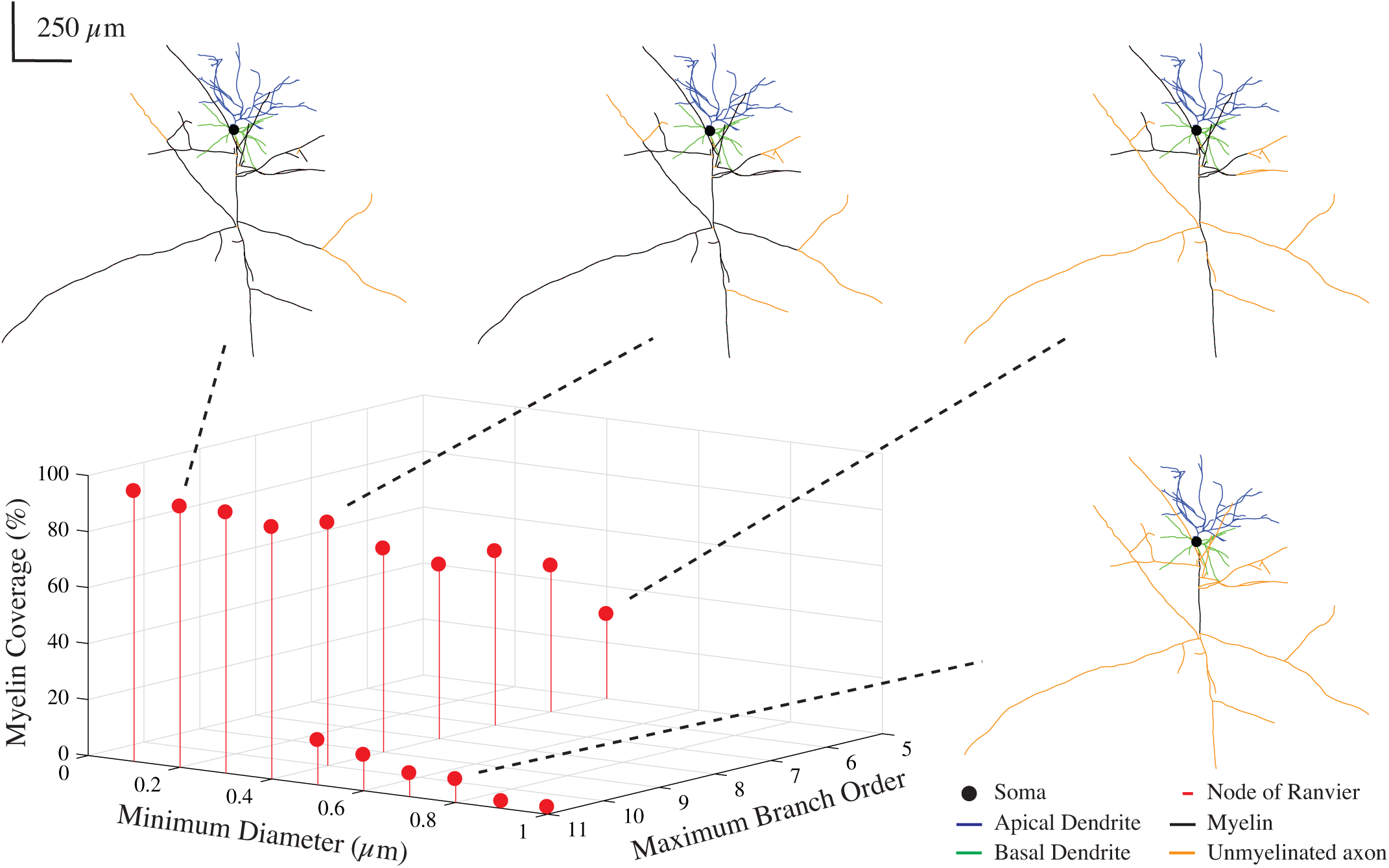
Method for scaling myelin coverage. Percent myelin coverage (by surface area) for example L2/3 pyramidal cell model as the minimum myelinated axon diameter and maximum branch order are varied. These two parameters were controlled to generate the patterns of myelination used in Fig. 6, which includes all unique values of percent myelin coverage shown here.

**Figure S3.**
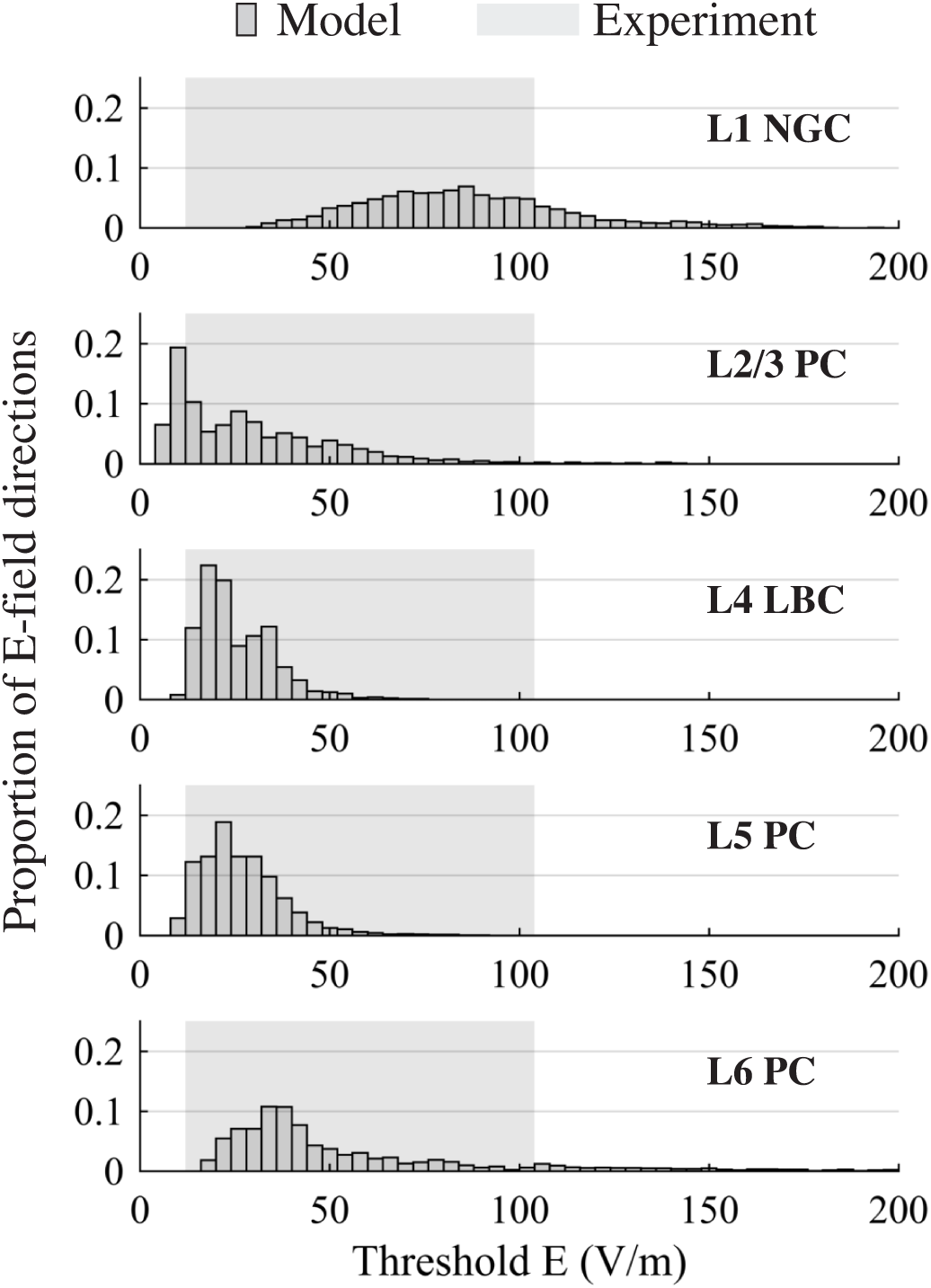
Thresholds for 100 ms, uniform E-field stimulation agree with *in vitro* experiments. Histograms of simulated threshold amplitudes for 100 ms, uniform E-field stimulation across all E-field directions for each cell type (5 clones per cell type). Shaded region indicates range of minimum E-field intensities at which excitatory post-synaptic potentials (EPSPs) were observed in [6].

